# Energy of functional brain states correlates with cognition in adolescent-onset schizophrenia and healthy persons

**DOI:** 10.1101/2023.11.06.565753

**Authors:** Nicholas Theis, Jyotika Bahuguna, Jonathan E. Rubin, Shubha Sankar Banerjee, Brendan Muldoon, Konasale M. Prasad

## Abstract

Adolescent-onset schizophrenia (AOS) is rare, under-studied, and associated with more severe cognitive impairments and poorer outcomes than adult-onset schizophrenia. Neuroimaging has shown altered regional activations (first-order effects) and functional connectivity (second-order effects) in AOS compared to controls. The pairwise maximum entropy model (MEM) integrates first- and second-order factors into a single quantity called energy, which is inversely related to probability of occurrence of brain activity patterns. We take a combinatorial approach to study multiple brain-wide MEMs of task-associated components; hundreds of independent MEMs for various sub-systems are fit to 7 Tesla functional MRI scans. Acquisitions were collected from 23 AOS individuals and 53 healthy controls while performing the Penn Conditional Exclusion Test (PCET) for executive function, which is known to be impaired in AOS. Accuracy of PCET performance was significantly reduced among AOS compared to controls. A majority of the models showed significant negative correlation between PCET scores and the total energy attained over the fMRI. Across all instantiations, the AOS group was associated with significantly more frequent occurrence of states of higher energy, assessed with a mixed effects model. An example MEM instance was investigated further using energy landscapes, which visualize high and low energy states on a low-dimensional plane, and trajectory analysis, which quantify the evolution of brain states throughout this landscape. Both supported patient-control differences in the energy profiles. Severity of psychopathology was correlated positively with energy. The MEM’s integrated representation of energy in task-associated systems can help characterize pathophysiology of AOS, cognitive impairments, and psychopathology.

## 1. Introduction

Adolescent onset schizophrenia (AOS) is associated with more prominent developmental and premorbid abnormalities with more severe cognitive impairments (Frangou, 2010; Holtmaat & Svoboda, 2009; Kester et al., 2006; Rapoport & Gogtay, 2011; Thaden et al., 2006), especially in working memory (Brickman et al., 2004; Karatekin, Bingham, & White, 2009; Karatekin, White, & Bingham, 2008; White, Mous, & Karatekin, 2013), executive functions (Frangou, 2010; Kester et al., 2006; Rapoport et al., 1997; Thaden et al., 2006), and attention (Oie & Hugdahl, 2008; Oie & Rund, 1999; Oie, Sundet, & Rund, 2010; Thaden et al., 2006), and with poorer long-term outcomes (Frangou, 2010; Kumra & Charles Schulz, 2008; Kumra, Shaw, Merka, Nakayama, & Augustin, 2001; Rapoport et al., 1997). Hence, AOS is proposed as a more severe form of schizophrenia (Kumra & Charles Schulz, 2008) although it is phenomenologically continuous with adult-onset schizophrenia (Frangou, 2010; Kumra et al., 2001; Rapoport et al., 1997). The neurobiology of AOS is under-investigated, evidenced by proportionately fewer peer-reviewed publications on AOS compared to adult-onset schizophrenia. Despite receiving greater attention recently, research on AOS constituted approximately 3% of all investigations on schizophrenia in peer-reviewed publications in each of the last 5 years **(Supplemental figure 1)**. In addition, although AOS constitutes 12.3% of all schizophrenia (Solmi et al., 2022), the economic and human cost of AOS far exceeds its proportional prevalence (Cloutier et al., 2016). Cognitive deficits are consistently associated with poor long-term outcome (Bora, Gokcen, Kayahan, & Veznedaroglu, 2008; Frith, 1996; Green, 1996; Velligan & Miller, 1999) and respond minimally to antipsychotics (Mishara & Goldberg, 2004; Woodward, Purdon, Meltzer, & Zald, 2005). Advanced methods could enable a better understanding of the pathophysiology of AOS, and lead to new treatment strategies with improved outcomes.

Most fMRI studies have examined group differences in the activation of brain regions, namely first-order properties (Gitelman, Ashburner, Friston, Tyler, & Price, 2001), or pairwise correlations of spatially distributed regions and derived functional networks based on co-activations, namely second-order properties (Friston, Holmes, et al., 1994; Friston, Jezzard, & Turner, 1994; Worsley, Evans, Marrett, & Neelin, 1992). These models provide limited information at the systems level. First-order models do not reveal spatiotemporal relationships in the evolution of the hemodynamic response. Second-order models rely on correlation between node pairs with the assumption that pairwise node correlations are independent of other pairs, and often do not provide mechanistic insights. Neither model adequately captures the temporal evolution of neural dynamics as a collective process.

The pairwise maximum entropy model (MEM) (Schneidman, 2016; Yeh et al., 2010) represents an underexplored but promising avenue of research. The MEM mathematically integrates both first-order activity and second-order pairwise-activations across brain regions (Yeh et al., 2010) in a way that relates the empirically measured probabilities of occurrence of brain states, which are temporal snapshots of activity, to the underlying network structure and external activation. The MEM does not assume independence of pairwise correlation between nodes but instead captures the probabilities of different individual nodes being active, and pairs of nodes being synchronously active during discrete time samples, thus focusing on collective activity patterns across brain regions (Lamberti et al., 2022). The pairwise MEM is considered to be a powerful tool to bridge the gap between micro- and macro-scale activation structure (Fortel et al., 2022). The MEM is related to the Ising model (Kloucek et al., 2023), which was originally proposed in the context of statistical mechanics of magnetic dipoles. Similarly, brain regions can be thought of as ‘active’ or ’inactive’ at a given point in time, can continuously change their activation state, and interact according to an underlying network configuration to produce brain states of varying likelihood. The MEM framework associates an “energy” to each brain state, also called the “integrated energy” of a system configuration when applied to fMRI data (Das et al., 2014), which has some similarity to the concept of potential energy in physics, albeit unitless.

Adapting the MEM from its original physics context (Brush, 1967) for use in fMRI is relatively novel; versions of the MEM have been applied to examine neural dynamics of cognitive performance using fMRI (Watanabe et al., 2013) for around a decade. The MEM approach is especially useful to investigate statistics of the time-varying dynamics during cognitive tasks because it models BOLD responses at every repetition time (TR) of the fMRI acquisition (Kang, Jeong, Pae, & Park, 2021) providing arguably the best temporal resolution for fMRI data analysis compared to traditional second-order comparisons including dynamic functional connectivity. Further, investigation of brain networks using a pairwise MEM and associated energy landscape analysis (ELA) provides a useful representation of network organization (Watanabe, Masuda, Megumi, Kanai, & Rees, 2014) that can be compared between different tasks, conditions, or subject groups (Watanabe & Rees, 2017). A prior study found multiple local minima in resting fMRI suggesting that in rest conditions, the brain naturally reverts to one of a collection of preferred states (Ezaki, Fonseca Dos Reis, Watanabe, Sakaki, & Masuda, 2020). Experimentally or environmentally induced perturbations reduced the number of local minima considerably, while other effects depended on the specifics of the perturbation (Kang, Pae, & Park, 2017). Another resting fMRI study showed different characteristics of the energy landscape in the default mode and frontoparietal networks (Watanabe, Hirose, et al., 2014). Energy and cognitive performance may be related (Jeong et al., 2021), in that energy level may reflect the complexity of the mental work being done or the efficiency of strategies adopted to complete a cognitive task; for example, complex tasks or inefficient strategies may lead to high energy and hence low probability states that would not naturally arise in more typical, less complicated settings.

Task-fMRI can provide data on differences in functional and network characteristics that can be used as biomarkers and as treatment targets. Since prior studies have reported impaired executive function in AOS patients (Frangou, 2010; Rapoport et al., 1997; Thaden et al., 2006), we selected the Penn Conditional Exclusion Test (PCET) as an fMRI task to test executive function. PCET is a part of the Penn Computerized Neurocognitive Battery that has been validated in schizophrenia (Gur, Ragland, Moberg, Bilker, et al., 2001) and healthy (Gur, Ragland, Moberg, Turner, et al., 2001) subjects and used in our previous studies (Kuo et al., 2018; **Prasad** et al., 2010; Prasad, Muldoon, Theis, Iyengar, & Keshavan, 2023; Roalf et al., 2013). The PCET is related to the Wisconsin card sorting test. The performance on the PCET has been correlated with verbal fluency and work behavior such as cooperativeness, work quality, and general impressions including quality of work, verbal fluency, and executive function (Kurtz, Wexler, & Bell, 2004).

We applied the pairwise MEM to fMRI data obtained while participants were performing the PCET task. The relationship between MEM energy, and accuracy of in-scanner PCET, and severity of psychopathology were examined. We hypothesized that: 1) total energies of the states an individual experiences will be associated with that individual’s cognitive performance and differ by group, and 2) energy landscapes will consist of one or more ‘wells’, or local minima, representing attractor states, and that good performers on the PCET will spend more time in the wells of a given energy landscape. We hypothesized that the control group would have larger well areas and hence a greater fraction of empirically observed states with lower energy than the AOS group. The shallower wells in the AOS group will be associated with higher variability in the single trial trajectories as compared to the healthy controls.

## 2. Methods

### 2.1. Sample description

We enrolled 23 AOS from in- and out-patient clinics of the University of Pittsburgh Medical Center, Pittsburgh and 53 adolescent healthy controls from the community. Adolescent-onset was defined as schizophrenia subjects who had an onset of psychotic symptoms after puberty but before completing 18 years of age. Puberty was assessed using the Peterson Pubertal Developmental Scale and all subjects had to score ≥2 to be eligible for enrolment. Participants diagnosed with intellectual disability according to the DSM-IV, having history of substance use disorder in the last 3 months, head injury with significant loss of consciousness, tumors, encephalitis, and had suffered neonatal asphyxia were excluded. Subjects were administered the Structured Clinical Interview for DSM-IV (SCID-IV) and selected items on the Kiddie-Schedules for Assessment of Depression and Schizophrenia (K-SADS). Consensus diagnosis by experienced clinical investigators was made after reviewing all available clinical data including the charts. The University of Pittsburgh Institutional Review Board approved the study. Informed consent was obtained from all participants after providing a full description of the study including the risks and benefits. We administered Positive and Negative Syndrome Scale (PANSS) (Kay, Fiszbein, & Opler, 1987) to all subjects to assess the severity of psychopathology. PCET scores are from the in-scanner task performance. Antipsychotic administration was calculated using the antipsychotic dose-years (Andreasen, Pressler, Nopoulos, Miller, & Ho, 2010) which converts different antipsychotic medications into chlorpromazine equivalents and the duration of administration where 1 dose-year is equal to 100 mg chlorpromazine equivalent per day taken for one year. The antipsychotic dose-year provides a better metric for calculating the duration and dose than the total dose alone.

### 2.2. Image Acquisition

Imaging data were obtained on a 7 Tesla whole body scanner. T_1_-weighted MP2RAGE scans were acquired in the axial plane (348 slices, 0.55 mm thickness, TE=2.54ms, TR=6 seconds, in-plane voxel matrix size of 390×390), at 0.55 mm isotropic voxels resolution. Functional MRI echo-planar images were acquired in the axial plane (86 slabs, 1.5 mm thickness, TE=20ms, TR=3 seconds, in-plane voxel matrix size of 148×148), at 1.5 mm isotropic voxel resolution over, and 142 volumes over time were acquired. Field maps were acquired for magnetic susceptibility distortion correction with the same acquisition parameters as the fMRI in terms of matrix size and slab number, but with a longer TE (36.2ms) and TR (6 seconds).

### 2.3. PCET Task

Functional MRI data was collected while subjects performed the PCET task **(Figure 1)**. Our choice of PCET performance is supported by previous studies that relate executive function to workplace behavior (Kuo et al., 2018). These domains are impacted by schizophrenia. Each trial presented four shapes and participants selected the shape that did not belong. The sorting principles changed, with feedback to help subjects develop new strategies. Subjects were expected to select the correct stimulus with the minimum number of errors despite changing sorting principles. Such set shifting was considered to indicate cognitive flexibility, which is subsumed as executive function (Kurtz et al., 2004). The number of correct responses was counted as the measure of accuracy of performance, with 48 possible total correct, but the task is designed such that a perfect score is not possible without lucky guessing and anticipation of rule changes.

**Figure 1:**
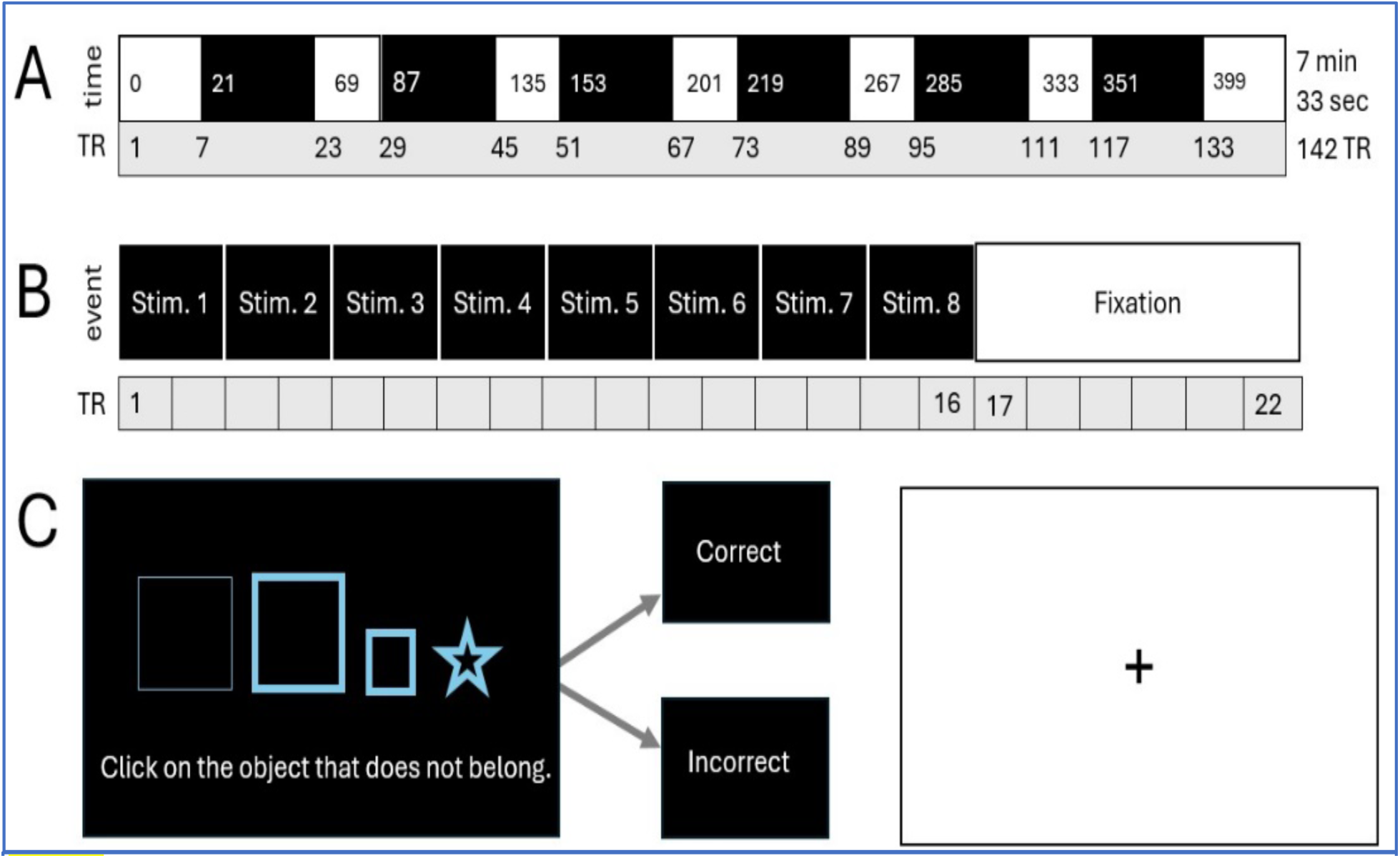
The Penn Conditional Exclusion Task (PCET) used in this study to collect task fMRI data related to executive functions. A) Six task blocks (black boxes) and 7 fixation blocks (white boxes) are shown. The onset time in seconds (numbers in white and black boxes) as well as the corresponding MRI TR (grey box) are shown. Each TR is 3 seconds with total number of TRs = 142; total fMRI duration 7 min and 33 sec. B) Within a task block, 8 stimuli are presented over 48 seconds followed by a fixation block consist of a cross. C) Each stimulus consists of 4 objects with varying features (line thickness, size, and shape). The participant’s responses are provided with feedback (“correct” or “incorrect”).

### 2.4. fMRI data pre-processing

All the pre-processing steps for the task fMRI data were performed using fMRIPrep 22.0.2 (Esteban et al., 2019), which included correction for motion (using `mcflirt`), slice time correction, susceptibility distortion correction, and brain extraction before registering to a standard space (MNI 152NLin6Asym) using the Human Connectome Project approach (Glasser et al., 2013). While more widely used for 3T imaging, fMRIPrep is also used for preprocessing of 7T fMRI (Miletic et al., 2022). Image registration of fMRI volumes to standard space was performed using co-registration with the T_1_-weighted image. Signal confounds were estimated, encompassing the global mean signal, and frame-wise displacement and DVARS (Power, Barnes, Snyder, Schlaggar, & Petersen, 2012; Power et al., 2014). Non-aggressive AROMA automatic denoising, with the default 200 components set by FSL MELODIC, was used to denoise each acquisition.

As the final step of pre-processing, all denoised 4D fMRI acquisitions were submitted to group independent component analysis (GICA), a data-driven strategy that defines spatial maps of voxel activity common across subjects (Varoquaux et al., 2010). This process reduces the high-dimensional fMRI into relevant population-level and likely task-driven BOLD signals that appear on the inter-subject level (Salman, Vergara, Damaraju, & Calhoun, 2019). GICA is performed using a larger selection of components than expected to be necessary to explain the task-associated regions. Because AROMA was already performed at the subject level, and because GICA is performed at the inter-subject level, the chances that individual GICA components are noise related is reduced. Here, we chose 70 components (Allen et al., 2014), which aims to have sufficiently many components to create a “functional parcellation”, with the understanding that too few components (<20) will result in fused functional parcels, and too many components (>100) will cause reproducibility issues (Abou-Elseoud et al., 2010). Where GICA resulted in a single, spatially non-contiguous component, that component was split into individual components.

### 2.5. Fitting the MEM and the Energy Model

A combinatorial design was used to assess statistics of interest across multiple models. This is because the MEM framework is limited by the exponential relationship between the number of nodes included in the model, N, and the resulting number of possible binary states for which energy will be calculated, B, where B = 2^N^. Modern anatomical cortical atlases have hundreds of nodes, for instance 360 in the popular Glasser atlas used in the Human Connectome Project (Glasser et al., 2016), resulting in 2.3×10^108^ unique binary states, which far exceeds the number of states that can actually be observed during our imaging data acquisition, and there is limited *a priori* information about which nodes are relevant for the task. Rather than attempting to establish localized cortical associations in an atlas, this analysis aims to find signal patterns that are common at the population level, then sample different combinations of subnetworks composed of these signals to assess features of the MEM common across many highly relevant systems.

To achieve this, the fMRI acquisitions were reduced to a relatively small number of task-associated components extracted from preprocessed fMRI using GICA (Varoquaux et al., 2010). Specifically in our analysis, in each of 501 samples (**Figure 2**), a selection of nine random GICA components was modeled as a nine-node MEM. Then, groupwise comparisons of energy between AOS and HC were examined followed by correlation between the total energy attained over the fMRI and PCET performance per subject assessed across all model instantiations.

**Figure 2:**
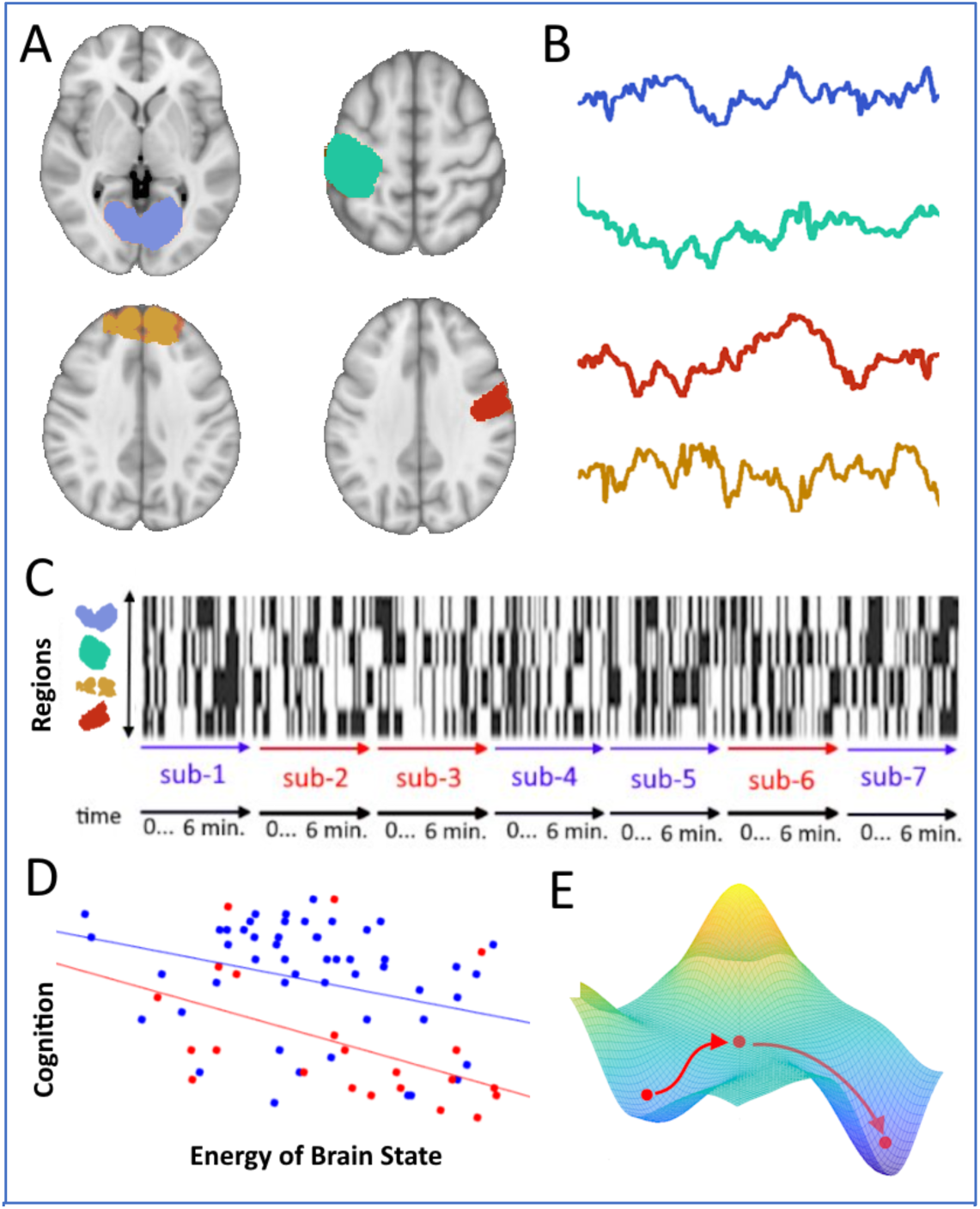
A schematic outline of the experimental design. One sample of nodes is pictured, with only four nodes. A. Four example components from GICA are randomly selected. B. An example time series for these nodes for a single subject. C. The four regions are binarized and concatenated across all subjects. Next, an MEM is fit to the data, using the binarized components as the nodal timeseries used to optimize the model parameters. D. The MEM assigns energy values to each brain state, and these values are summed per individual and compared performance on the PCET. The relationship between total energy and cognitive performance across subjects is the main variable of interest compared across node samples. E. For one sample, an energy landscape analysis is performed to study differences between the group-level trajectories on the population level MEM.

The resulting components from GICA are z-scored over the entire task, and thus continuously valued, but MEM input data must be binary. To prepare the z-scored component data, a simple threshold of zero was selected for binarization (Fortel et al., 2022), rather than a threshold with respect to the fixation period which was too short in our data for the signal to reach a baseline. Additionally, the number of GICA components is expected to be around ten times greater than the acceptable system size for the MEM. To choose components in a robust and unbiased way, 501 samples were selected, where each contained 9 randomly selected components. The timepoints for all subjects were concatenated together in a matrix, *T*, with *N* rows representing regions and a number of columns, *M*, equal to the total number of timepoints (142 per subject) in the entire sample across all subjects (n=76), i.e. 10,792 observations per instantiation The threshold choice for binarization can affect the outcome of the energy model **(Supplemental figure 2)**. The seven threshold choices tested for binarizing the BOLD-derived observations, z-scored across all subjects and timepoints, were 0, +/-0.5, +/-1.0, and +/-2.0. For example, a threshold of 0 would produce a matrix *T* for which roughly half of all elements took the value 1, with correspondingly fewer or more 1 values for higher or lower thresholds, respectively. Our choice of the threshold 0 for our subsequent analyses was justified by the observations from the application of different thresholds because it produced the most informative energy landscape with the best fit of the energy function to the empirical probabilities of state occurrence; additionally, this threshold choice has been used in previous studies on the MEM for fMRI (Fortel et al., 2022).

### 2.6. Energy Model

The binary state matrix (**Figure 2C**) contains many repeating entries due to the underlying probability distribution of occurrence of the binarized activation states. This matrix is distinct from the matrix of all possible state vectors, each represented as *V_k_*, where the index *k* runs from 1 to *B=2^N^*, which is 512 in the case of N=9. Each *V_k_* represents a binary configuration, i.e., an “on/off” pattern of nodal activity. Energy, *E*, is defined for each possible state, *V_k_*, according to the pairwise MEM (Yeh et al., 2010) as:

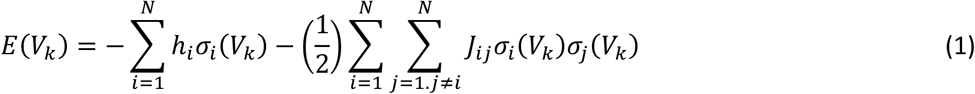

where the *N*-by-*N* matrix *J* and the *N*-vector *h* are parameters to be determined from the imaging data using a fitting algorithm **(Supplemental Material 3)**. In equation (1), each term σ_*i*_(*V_k_*) takes the value 1 if region *i* exhibits suprathreshold activation in state *V_k_* and takes the value 0 otherwise. While the energy value itself has no absolute interpretation and only relative changes in energy are meaningful, the energy of a state maps to the probability of a state, *P(V_k_)*, as follows:

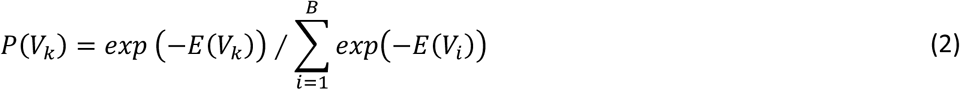

We evaluated the fit of the model by comparing the estimated and actual first-order (mean) and second-order (covariance) features of the data **(Supplemental Figure 3)**. As an additional check, we examined the relationship between the energy and probability of occurrence across all states as estimated by the model and the empirical probability of each state in the data, which also demonstrated a good fit (**Supplemental Figure 4**).

### 2.7. Statistical Comparisons: Cognitive, Psychopathological and Clinical

From equation (1), we derived a vector of energy values along a sequence of time points for each subject, according to the respective group level MEM for each sample of node combinations. A two-sample t-test was used to compare the distribution of PCET scores by group. A mixed effects linear model taking into account repeated energy measurements from each individual was considered to estimate the group differences of energy level across all samples. Pearson’s correlations between individual PCET scores and total energy over all time were calculated for the whole group and AOS and controls groups separately. This comparison was performed for all node samples. Pearson correlation tests were also used to examine the relationship of energy of each subject with severity of psychopathology assessed using the PANSS in the example model. First, the correlation of total PANSS score in relation to total energy obtained from the MEM was examined. Next, an examination of PANSS scores for each group of symptoms, namely total positive, total negative, total disorganization, and total cognitive score was made. The correlation of PCET accuracy scores with the severity of psychopathology was also tested. The results are reported after correcting for multiple tests using the Bonferroni correction.

### 2.8. Energy Landscape via Projection onto Principal Components

Because of the challenges of visualizing hundreds of models, energy landscape analysis was only performed on a single 9-node example model that was selected at random from among the models where a significant correlation between PCET performance and energy was detected, to help understand what features are being observed throughout the MEM. Since the basis for the energy value is the *9*-dimensional brain state configuration across the nodes, and time points with the same energy value could correspond to very different brain states, we used Principal Component Analysis (PCA), performed on *T*, the matrix of observed binary states concatenated across all subjects and time points, to describe states in terms of a lower-dimensional manifold.

The PCA isolates the orthogonal eigenvectors of the covariance matrix, or principal components (PCs), that explain the most variance in the data. Our aim was to visualize an energy landscape in a three-dimensional state space, where two dimensions represent the data projected onto the first 2 PCs and the 3^rd^ dimension represents the energy corresponding to the state. In this space, moving along a PC axis changes the weight on that PC vector; different states are approximated by different weighted sums of the PC vectors and thus different states are projected to corresponding points in PC space. The energy landscape was smoothed with a 2-dimensional Gaussian kernel, with σ=1.5; in brief, the smoothing process introduces a kernel centered at each data point and computes a smooth function by summing over these kernels. The smoothed version was used to visualize the energy manifolds, but all quantification was done on the individual states of the MEM. The same value of σ was used to generate all energy landscapes. The basins of attraction of energy minima correspond to low (usually negative) energy values represented with cooler colors, whereas the peaks correspond to higher (usually positive) energy values represented with warmer colors.

### 2.9. Energy Landscape Analysis (ELA) and Energy Landscape Trajectories

We characterized the difference between the energy landscapes of the control and AOS groups using separate groupwise ELAs from the example model’s MEM. The ELAs were assessed for “well” features, i.e. basins or sinks that represent collections of high probability states. A well is defined by all states in the neighborhood of a local energy minimum with energies that fall below a chosen threshold, also called a well boundary. To perform the comparison, we used the cumulative distribution function (CDF).

The 6 task blocks of 22 TRs (3 secs each) of the PCET were averaged for each participant to obtain a corresponding energy trajectory. The trajectories were smoothed using a box car kernel of length 5. The top 5 performers from the control and poorest performers on the AOS group were chosen on the basis of the 5 highest or lowest PCET scores, in order to achieve the highest contrast between the groups, and to lower variance within group level trajectories. The states for these selected subjects were projected onto the PC dimensions and smoothed to derive their energy landscapes. An ellipsoid for each group was calculated by finding the average standard deviation across X (PC1), Y (PC2) and Z (Energy) axes corresponding to the AOS and control trajectories, respectively. The difference between the distributions of pairwise average trajectory distances was measured with a two-sided T-test.

## 3. Results

### 3.1 Demographics and clinical characteristics

HC and AOS differed significantly in sex distribution but not in age. Healthy controls showed significantly higher accuracy on the PCET task compared to the AOS group **(Table 1**; **Figure 3).** Irrespective of clinical group, the males (N=30) scored on average 31, while the females (N=46) scored on average 35, but the difference was not significant in a two-sample t-test (p=0.083, t=-1.7382). Mean antipsychotic dose-years was 2.69±6.77 meaning the AOS patients were on 269 ± 677 mg chlorpromazine equivalent mg per day per year. Medication compliance was highly variable that may explain a large standard deviation.

**Table 1:** Demographic and clinical characteristics

|  | EOS | HC | Statistics |
| --- | --- | --- | --- |
| Age (years) | 19.83 (1.47) | 19.64 (1.64) | $t=0.47$, df 74, $p=0.64$ |
| Sex |  |  |  |
| Male | 8 | 17 | $\chi^2=5.51$, df 1, $p=0.02$ |
| Female | 15 | 26 |  |
| Duration of illness (years) | 1.43 (2.70) | 0 | -- |
| PANSS score* |  |  |  |
| Total | 59.52 (14.68) | 31.13 (2.51) | $F(1,76)=147.07$, $p<0.001$ |
| Positive | 16.96 (4.52) | 7.17 (0.51) | $F(1,76)=184.08$, $p<0.001$ |
| Negative | 12.96 (4.44) | 7.57 (1.67) | $F(1,76)=40.96$, $p<0.001$ |
| General | 29.61 (7.50) | 16.40 (1.35) | $F(1,76)=132.65$, $p<0.001$ |
| PCET Total score | 26.43 (8.89) | 35.08 (8.42) | $F(1,76)=3.98$, $p=0.05^{\#}$ |
| WASI Total | 100.78 (12.23) | 107.59 (11.23) | $F(1,76)=1.27$, $p=0.26^{\#}$ |

\*MANCOVA model consisting of PANSS component scores covarying for age, sex, and medications, Wilk's $\lambda=0.263$ , $F=64.50$ , $p<0.001$ ; $^{\#}$ Covarying for age, sex, and medications

**Figure 3:**
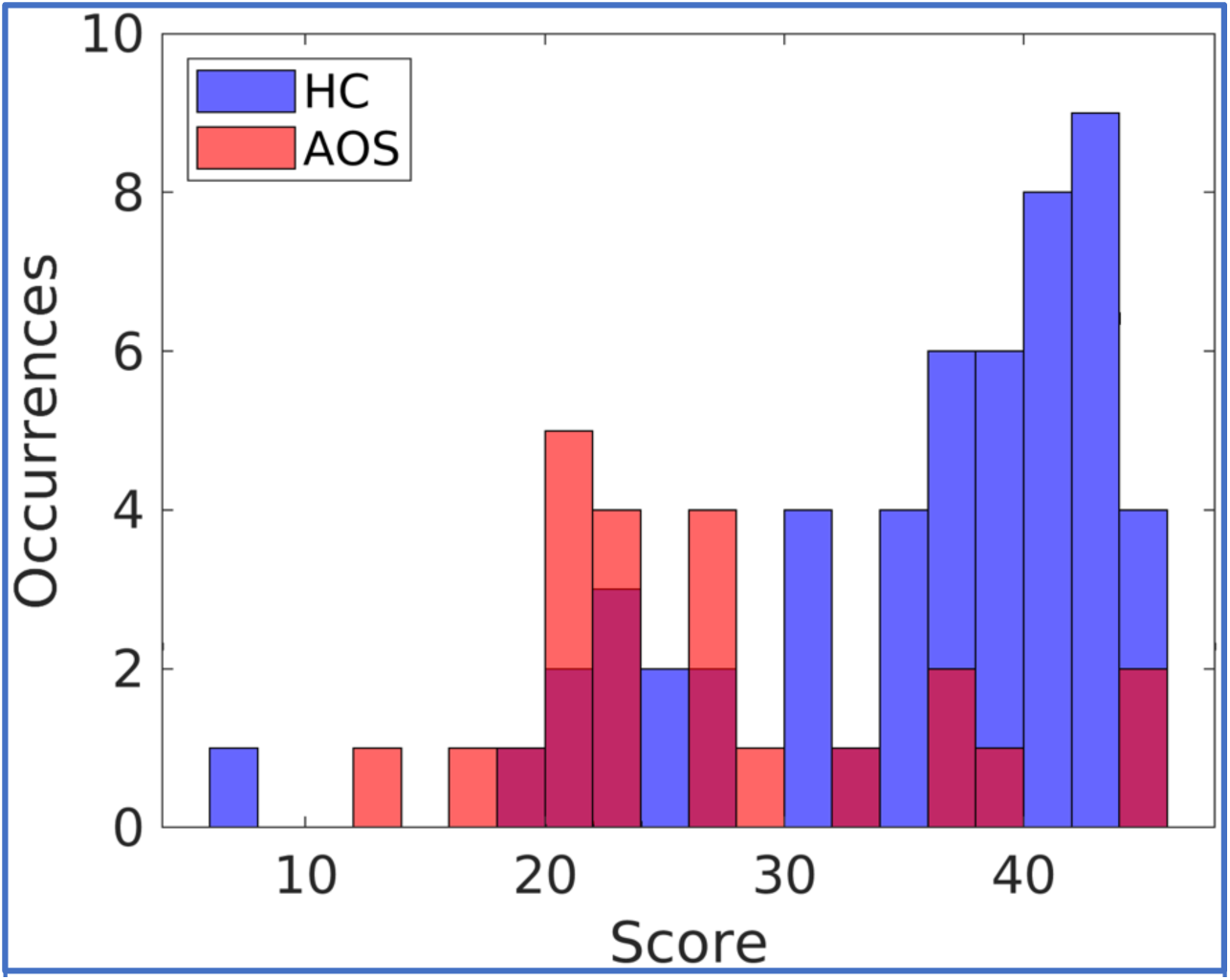
PCET scores distribution between AOS and HC. Controls show a right shift in higher PCET accuracy scores compared to patients who mostly concentrated to the left. Blue: controls. Red: AOS. X-axis is total PCET correct. Y-axis is the number of participants in the group who scored the corresponding amount.

### 3.2. Total energy across all permutations

In total, 92 spatially non-contiguous GICA components were identified, and these components were randomly sampled to obtain 501 sub-networks of 9-nodes each. The MEM for each sample resulted in a population-wide model that defines the energy of each state according to (**Equation 1**). This yields a 9 node by 142 timepoint matrix of energy values over time for each of the 76 subjects. The minimum energy observed among all states and model instantiations was –9.55, and the maximum 100, with a mean state energy of -1.36. A mixed effects model indicated the AOS group had significantly (p < 0.0001) higher energy by 0.031 units with a standard error of 0.00033. The model accounted for the correlations introduced by the repeated measures of the individuals among s, with the variability of the mean energy measurements for the different individuals accounting for around 73% of the total observed energy variability.

Among 501 subnetworks, 359 correlations of energy with PCET score were significant in the combined sample of AOS and HC (p<0.05). When the HC and AOS groups were considered separately, we observed significant correlations in 97 samples among controls and 180 among the AOS. These results are summarized in **Figure 4**, where the top row indicates the correlation value, *r*, the bottom row indicates the significance (-log of p value), with values further to the right being more significant and the vertical red line indicating the alpha value of 0.05. The yellow bin indicates the bin containing an example model on which we focused for further analysis.

**Figure 4:**
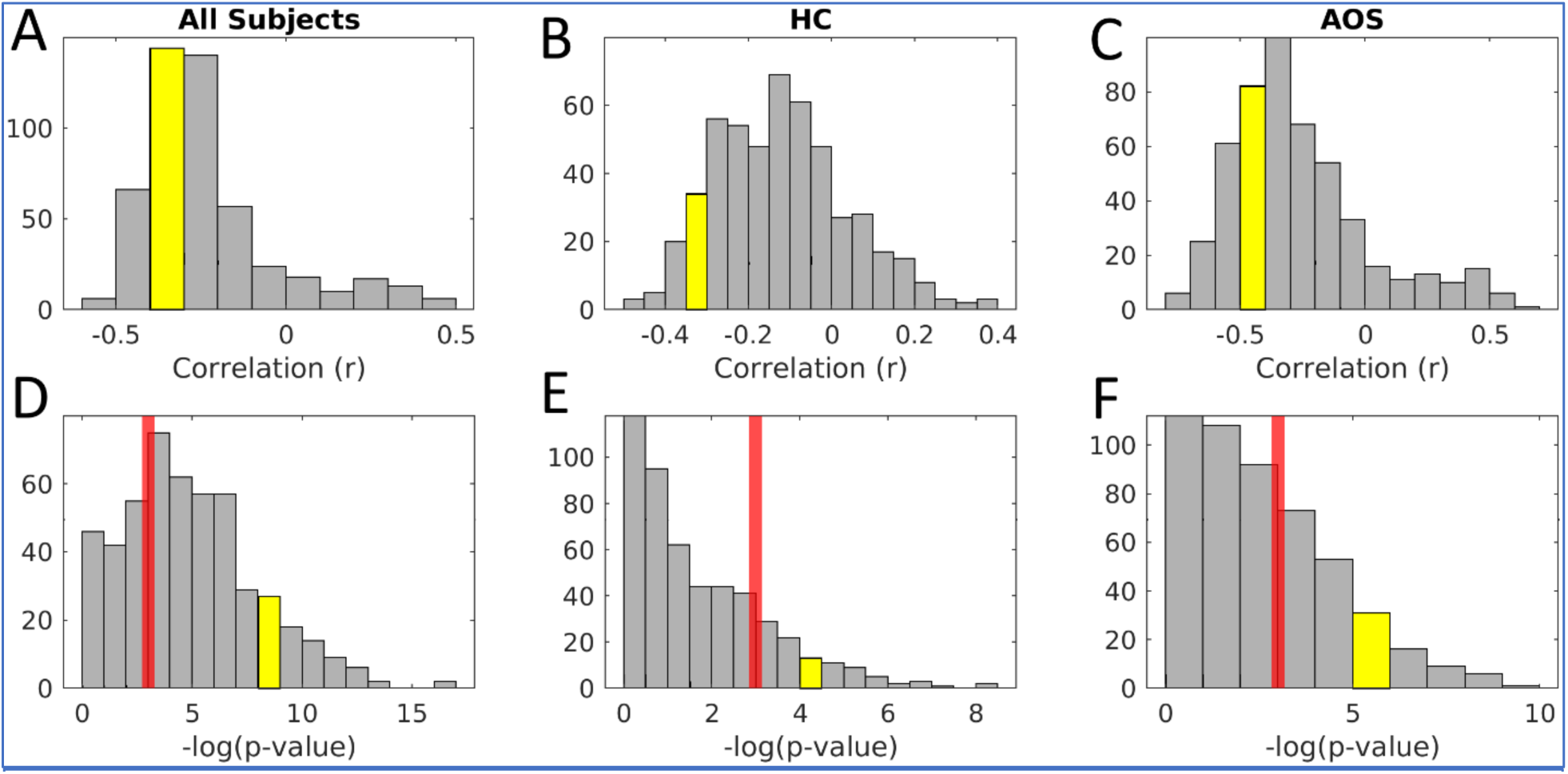
Histogram showing results of 501 sampling experiments for correlation between energy and PCET score. The yellow bin in each plot indicates the bin that the example model of interest belongs in. Plots in the top row show Pearson correlation, *r*, across subjects between the total energy that subject achieved over time and that PCET score. Plots in the bottom row show the significance, *p*, where the red vertical line indicates an α <0.05. Bins further to the right on the bottom plots are more significant due to the negative log operation. The left column shows group level results, the middle column shows results for the HC group, and the right column shows the AOS group results.

### 3.3. Total Energy in the Example Model

In the example model, the groupwise time-series visually suggests that the energy was higher in AOS during most of the task blocks compared to HC (**Figure 5A**). The total energy attained by each subject over all 142 fMRI timepoints is plotted against the PCET scores for each subject in the example model (**Figure 5B)**, and significantly correlated in the combined sample (N=76, r=– 0.39, p=0.0004) and separately for the control group (N=53, r=–0.30, p=0.03) and AOS (N=23, r=–0.48, p=0.019). Total energy also correlated positively with total PANSS score (r=0.23, p=0.04). Correlation with severity of psychopathology of each domain (positive, negative, and general psychopathology) were not significant after Bonferroni correction. The PCET score correlated negatively with total PANSS total score (r=-0.35, p=0.002), total positive (r=-0.38, p<0.001) and general psychopathology (r=-0.33, p=0.003) scores that survived the Bonferroni correction.

**Figure 5:**
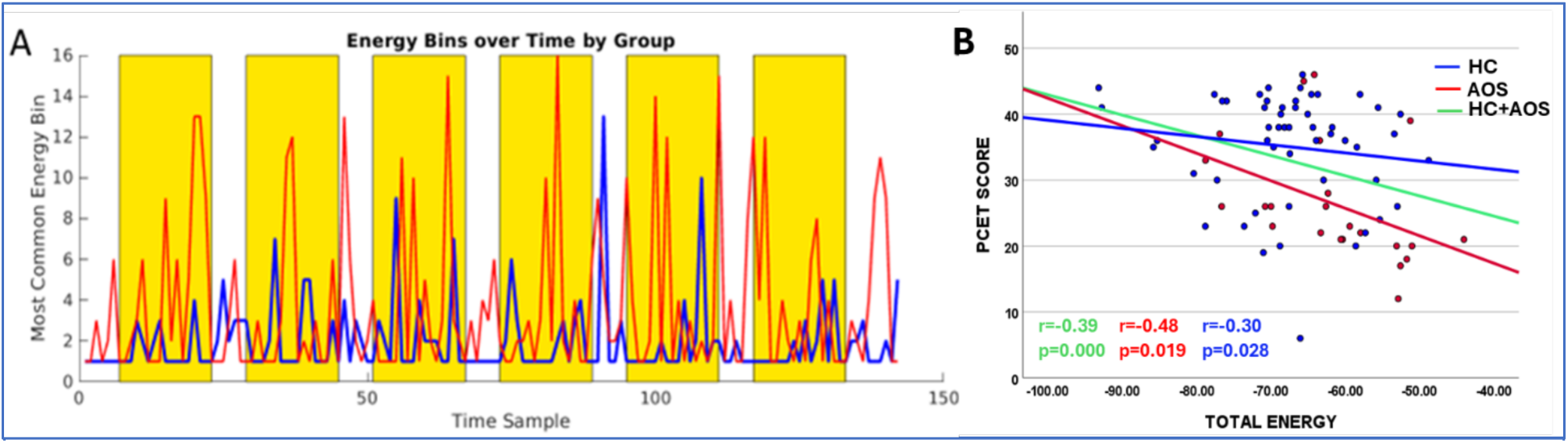
Energy Characteristics by Group for the single example sample of nodes. A. The energy level of the most frequent (modal) energy bin in each subgroup: blue is the control group (N=53), and red is the AOS group (N=23). Time samples are fMRI TR, which are 3 seconds in duration. The yellow regions represent the task blocks, which contain 8 stimuli each. Stimuli are presented every 9 seconds. B. Average Energy versus PCET Score. The energy values averaged across the entire duration of the fMRI (142 timepoints) for each subject (Blue point: healthy controls; Red points: AOS; Green: Combined sample of AOS and HC) show significant negative correlation with PCET scores.

### 3.4. Energy Landscape Analysis highlights group level differences

We characterized the difference between the energy landscapes of the control and AOS groups using separate groupwise ELAs from the example model’s MEM. Averaging of the energy levels across groups as well as separately for each group yielded energy landscapes with two dominant wells, which appeared to be shallower for the AOS group (**Figure 6A**). The cumulative distribution functions (CDF) for energies of states in each group (**Figure 6B**) showed a dearth of low-energy, high-probability states for the AOS group. In theory, any sufficiently low energy level could be taken as a threshold to define well boundaries, and well areas could be computed based on all states with energies below that level. An example for a well threshold of -1.2 is shown as green rectangle and the inset shows the calculation of the area under the CDF curve (AUC) up to this threshold for both the groups (**Figure 6B**). The AUC for each group quantifies the fraction of states that lie in the well (i.e., below that well threshold); the higher the well area, the higher the density of states with energy levels lower than the well threshold.

**Figure 6:**
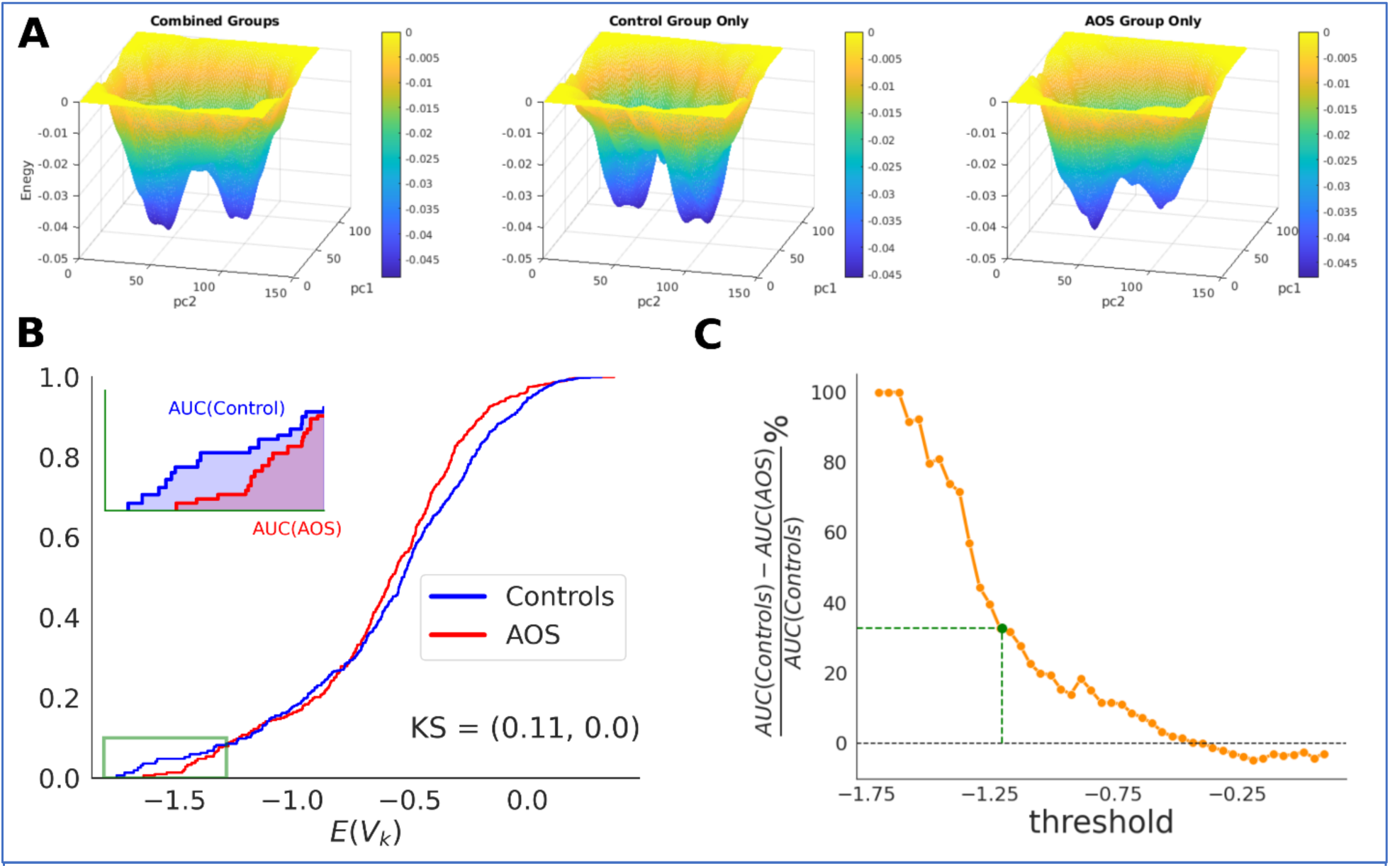
Energy landscapes for both groups show distinct wells. A) Comparison of energy landscapes for combined model (left) versus separate models for controls (middle) and AOS patients (right). All three manifolds show two primary wells, although the well depths and energy barrier between wells differ across them. Wells (blue regions) represent more probable, lower energy conglomerates of states. Theoretically, an individual’s brain activation states yield a sequence of positions on the energy landscape during cognition. B) Comparison of the well areas between MEM of controls versus AOS. The area under the curve (AUC) was calculated for energy cumulative distribution functions (CDF). The definition of well depends on the choice of a threshold. An example for threshold = -1.2 is marked with green rectangle, i.e. all states that have energy values lower than -1.2 are considered as well and the area under the curve (shown as an inset) calculates how many states form a well . C) Percent difference in AUC of control vs AUC for AOS group for different values of threshold. The total well AUC for control was 32.94% higher than for the AOS group as marked by green point and green rectangle in (B).As can be observed, the total well area in the for the control group is higher than the well area for AOS group for all values of threshold < -0.35.

To test our hypothesis that the healthy controls have larger AUCs, we compared the two groups by calculating the percent difference between the AUCs of controls and AOS. Since the difference between the well areas depends on the choice of threshold or the well boundary, we compared the well areas for healthy vs AOS patients for 45 thresholds between the range (−1.7, 0.1) as shown in **Figure 6C**. For a threshold of -1.2, for example, the well area for control group was 32.94% higher than well areas for the AOS patients (shown as the green point on the curve). It can also be observed that the well area for the control group is less than for the AOS group for all choices of thresholds less than -0.35 (**Figure 6C**), suggesting that although the comparison of well areas is dependent on the choice of threshold to define a well, it is extremely robust to this choice. This result aligns with our previous analysis showing that controls experienced more states in lower energy bins compared to AOS patients.

### 3.5 Shallower energy wells and higher energy in AOS correspond to more variable single trial trajectories that correlate with higher cognitive scores

To achieve the highest contrast between the groups, we selected the top 5 performers from the control group and the bottom 5 performers among AOS on the PCET task for ELA trajectory comparison. The trial trajectories (including the brief fixation period and the task block) represent averages of 6 such blocks that a subject visited. The individual trajectories for 5 control and 5 AOS subjects are shown as thin lines of cool and warm colors respectively (**Figure 7A**). As can be observed, the trial onsets always corresponded to high energy levels, which switched to lowest energy levels at stimulus onset. At the end of the trials, the dynamics return to high energy levels. The AOS trial trajectories traverse higher energy levels as compared to controls, as expected from their respective energy landscapes (**Figure 6A**), supported by the observation that the lowest functioning AOS also traversed longer distances on average (t-stat=-4.4, p<0.0001, **Figure 7B**). The AOS trial trajectories also show more variability as depicted by the ellipsoid that encompasses the average standard deviation of the spread of trajectories in X (PC1) and Y (PC2) space (**Figure 7C**).

**Figure 7:**
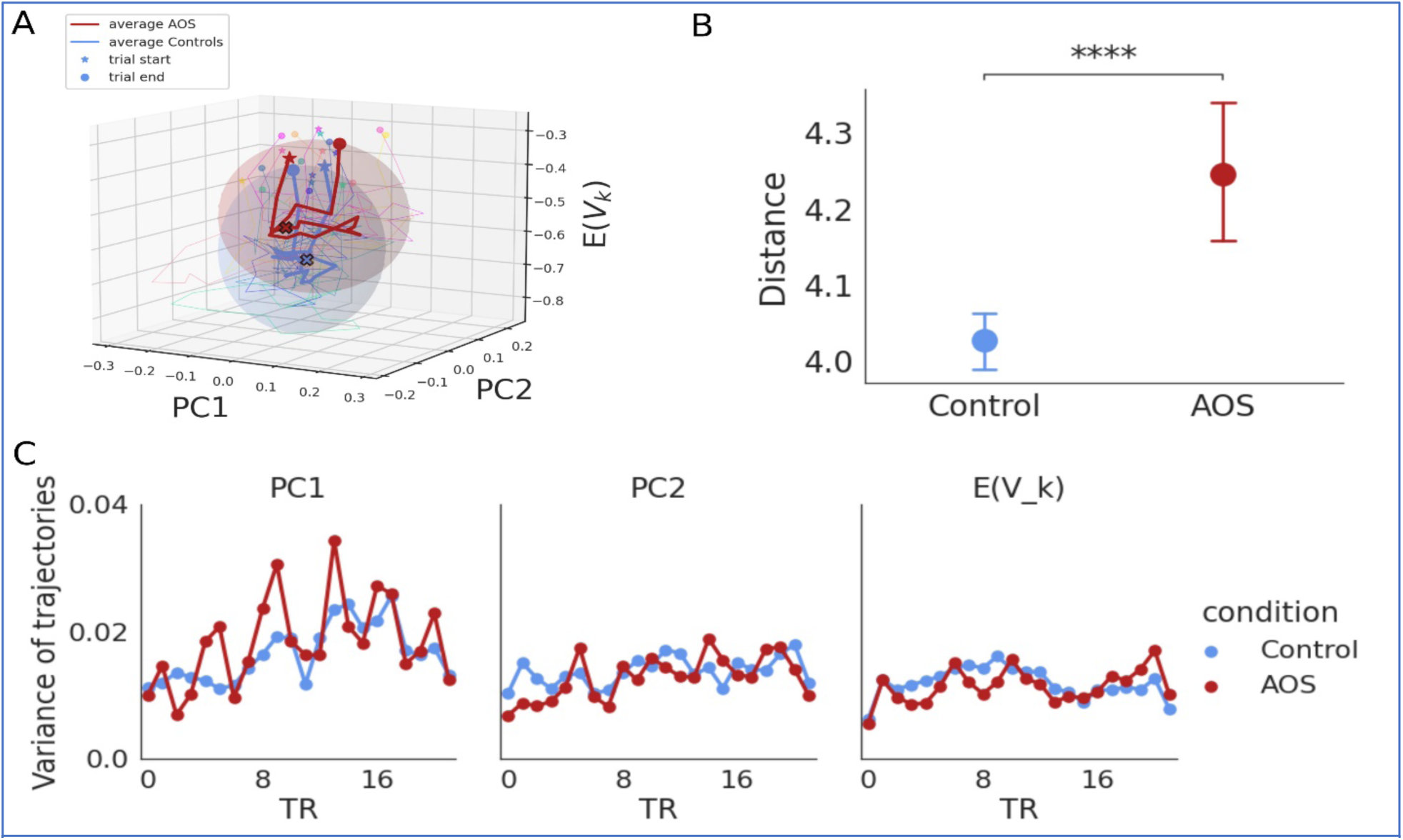
Average single trial trajectories for AOS have more variance compared to Controls: **A)** Individual average control trajectories for 5 controls are shown in cooler colors and thin lines (blue to green) and 5 individual AOS trajectories are shown in warmer colors (pink to yellow). The star (*) represents the start of the trial and the circle marks the end of a trial. The average trajectory over control individuals and AOS individuals are shown as thick blue lines for controls and as red lines for AOS. The filled X represent the stimulus onset during the trial. The ellipsoid encompasses the average standard deviation across all controls or AOS subject trajectories in X (PC1), Y(PC2) and Z (energy) axes. A higher radius of the ellipsoid along X and Y axis as seen for AOS suggests higher spread for AOS trajectories over PC1 and PC2. **B)** The pairwise Euclidean distances between the trajectories for Control group vs AOS group. The distances in Control group were significantly lower than the distances in the AOS group (4.03 vs 4.24, two sided t-test: -4.4, p= 0.0). **C)** Variance across the trajectories plotted as a function of TR for the three dimensions, i.e. PC1, PC2 and Energy. The trajectories in PC1 show the highest variance.

## 4. Discussion

We applied, for the first time, a pairwise MEM to understand the characteristics of energy, an integrated measure of brain regional activation and coactivation, among AOS and healthy subjects. The primary aim of this study was to determine if there is a relationship between energy, executive function, and psychopathology as well as diagnostic groupwise differences. A mixed effects linear model demonstrated that energy is significantly different between HC and AOS; a small but detectable increase of energy was associated with the AOS group. Across sampling of nodes, total energy across all states for a given subject over time correlated negatively with executive function (PCET score) in most sampling of nodes (**Figure 4**). While this relationship was more prominent at the whole-population level, likely because of the larger sample size, it was also observed within patient and control groups. Together, these findings suggest that a detectable, and unstable activation-coactivation pattern may be a potential biomarker of AOS, unfavorable executive function performance and greater severity of psychopathology.

Another aim of the study was to identify what energy landscape and trajectory features are associated with these characteristics. An example model (**Figure 5**) of interest was investigated, where significant differences in the groupwise energy landscapes were observed (**Figures 6**) and the AOS landscape had shallower wells. Differences in the average trajectories of the highest performing controls versus the lowest performing patients were also observed in the example model **(Figure 7**). Energy also positively correlated with severity of psychopathology measured using the PANSS in the example model, across all subjects. Greater severity of psychopathology also correlated negatively with performance on PCET across the whole population of both patients and controls. Thus, the observed relationship among energy, executive function performance, and severity of psychopathology suggests that high energy states may be relevant to pathophysiology.

Importantly, energy in the MEM is not a physical quantity but rather corresponds to the frequency with which states occur. The prevalence of high energy states in the AOS group and its association with lower PCET scores may be suggestive of inefficient or inconsistent processing during task performance. Overall, our findings suggest that the application of MEM to examine functional brain network configuration is a useful strategy especially since improvement in cognitive and social functions following certain treatments can be tracked using longitudinal functional imaging (Keshavan, Eack, Prasad, Haller, & Cho, 2017). A separate study by our group (under review) found that high energy states may be associated with schizophrenia even among adults and in the resting state, suggesting the MEM energy measure may be a distinct marker of schizophrenia relative to bipolar and major depressive disorders that showed different patterns of changes (Theis et al., 2024). Independent replication of these findings can enhance our understanding of the dynamically changing neurobiology that occurs during cognitive processing.

The utility of the pairwise MEM rests in its ability to potentially integrate findings from existing first- and second-order models in a manner that provides statistical insights into the network architecture, and activation patterns of complex systems, in detail. There are limitations to examining functional connectivity alone despite its strengths, e.g., functional connectivity implicitly assumes that the pairwise correlations are independent of each other and identifies two regions as ‘connected’ when they show correlation of BOLD signals (Adachi et al., 2012; Zuo et al., 2012). The MEM, however, is a conceptually different approach to infer global brain activity patterns that incorporates pairwise correlation of nodes where connection strengths are estimated in the context of the whole system, rather than independently for each node pair.

However, there are limitations to our study. The MEM places restrictions on the system size and dependence on the choice of threshold for binarization. In the case of system size, innovative methods of inverse Ising inference can help relax system size restrains, such as the pseudolikelihood estimate (Fortel et al., 2022). Binarization poses a difficult problem, because fMRI is unitless and un-calibrated but continuously valued (Bush & Cisler, 2013), which necessitates techniques like *z*-scoring, with the binarization threshold often depending on experimental conditions. We have attempted to address the restriction on system size by conducting repeated random sampling of the nodes, examining the results using robust statistical approaches, and reporting results after correcting for multiple testing. Furthermore, due to AOS being epidemiologically less frequent, even samples that are relatively large from the perspective of recruitment of AOS participants, as we have here, appear relatively small insofar as statistical power. Although neuromaturational processes might have affected some of the results, it is likely to be minimal because the mean age of both AOS and HC were in late adolescence and the difference in mean age between the two groups was not statistically significantly different. Thus, while encouraging, we emphasize the need for replications to rule out the role of potential confounds in our findings. Finally, while the strategy of using GICA components is useful to reduce data dimensionality and make the problem of modeling more tractable, ICA methods create regions of interest that do not necessarily respect anatomical boundaries to the extent that traditional brain atlases do, frustrating localization attempts (Petersen, Seitzman, Nelson, Wig, & Gordon, 2024). While useful as a data-driven method to detect population-level task-associated signal components, the spatial interpretation of GICA is less straightforward.

The association of energy with severity of psychopathology is intriguing and clinically important. Our study suggests that high energy states are correlated not only with impairments in executive functions but also with greater severity of psychopathology. Prior studies have shown that executive function deficits correlated with psychotic symptoms although more strongly with negative and disorganization symptoms than positive symptoms (Dibben, Rice, Laws, & McKenna, 2009; Dominguez Mde, Viechtbauer, Simons, van Os, & Krabbendam, 2009) and are also a major determinant of long-term outcome (Fett et al., 2011; Green, 1996). More research is needed to further elucidate precise mechanisms that link higher energy states with greater severity of psychopathology and executive function impairments in larger sample sizes.

Future research is needed to study full-brain MEMs, for instance using the 360-cortical node Glasser atlas (Glasser et al., 2016) and focus on elucidating trajectories of brain states (Transtrum & Qiu, 2016) through an energy landscape where wells represent higher probability states that correspond to attractors for the dynamics of neural activity. With more data, a future step would be to consider subject-level trajectories on the energy landscape defined by the sequence of activation states each subject exhibits during data collection, with possible clustering of trajectories to identify distinct phases and strategies of task performance.

## 5. Conclusion

Neuroimaging research into AOS has not previously considered the pairwise MEM framework, and we have presented evidence suggesting quantities from this model, particularly energy, are related to AOS, executive function, and psychopathology severity. The application of the MEM to fMRI has emerged in the last decade and represents an advancement over traditional fMRI analysis techniques, including correlational functional connectivity and regional activation modeling, because it provides a framework relating brain connectivity and temporal activity to the probability of observing system-wide activation patterns. In this study, MEM energy, which is inversely related to probability, was found to be associated with diagnostic groups, cognitive performance on the in-scanner PCET task, and total PANSS score. Patient-control differences in total energy over time were also observed across all node samplings, which results in altered energy landscapes, and temporal trajectories on these landscapes that are tied to groupwise differences in executive function. Schizophrenia symptoms in adolescent onset patients, therefore, may be associated with the presence of unfavorable, higher energy brain states during cognitive load.

## Data availability

The data will be made available through NIMH data repository.

## Acknowledgments

Authors thank Ms. Diana Dworakowski, Ms. Shaelyn Coles, Ms. Lydia Harvey, Mr. Dylan Tomsey, and Ms. Molly Stevens for their efforts in enrolling and characterizing participants. Authors also thank Dr. Matcheri Keshavan MD and Dr. Shaun Eack PhD for arriving at consensus diagnosis on participants after reviewing the clinical evaluations. We wish to thank Dr. Satish Iyengar, PhD Professor of Statistics, University of Pittsburgh, Pittsburgh, PA USA for his guidance for the statistical analysis.

## Author Contributions

NT performed analysis and drafted the manuscript; JB created the MEM implementation, performed analysis and participated in editing the manuscript; JR conceptualized the study design, supervised the analysis, and participated in drafting and editing manuscript; SB performed the statistical mixed effects model analysis; BM preprocessed the imaging data and ran GICA; KP managed the enrolment and characterization of participants in the study including obtaining the MRI data, conceptualized the study design, supervised the analysis, participated in drafting and editing manuscript, and in obtaining funds.

**Code availability** Python code for the gradient descent implementation is available on request.

## Supplementary Materials

### Supplemental material 1.

We surveyed publications of peer-reviewed manuscripts among topics including all schizophrenia, early onset schizophrenia, adolescent schizophrenia, schizophrenia over 19 years of age and schizophrenia from 1 month to 18 years of age on PubMed. The number of peer reviewed publications from 1921 until 2023 that we obtained from PubMed using the key words “schizophrenia”, “early onset schizophrenia”, and in addition using age as a criterion for each term (accessed on February 20, 2024). The ages that were used for retrieving the references were ≥19 years and between 1 month and 18 years. The data shows that only a fraction of peer-reviewed publications have been on early onset schizophrenia. This emphasizes the need for more research in this population. Further, the onset of psychotic symptoms during the period of adolescent brain growth can provide insights into how the maturation process and clinical manifestation are intertwined.

**Supplemental Figure 1:**
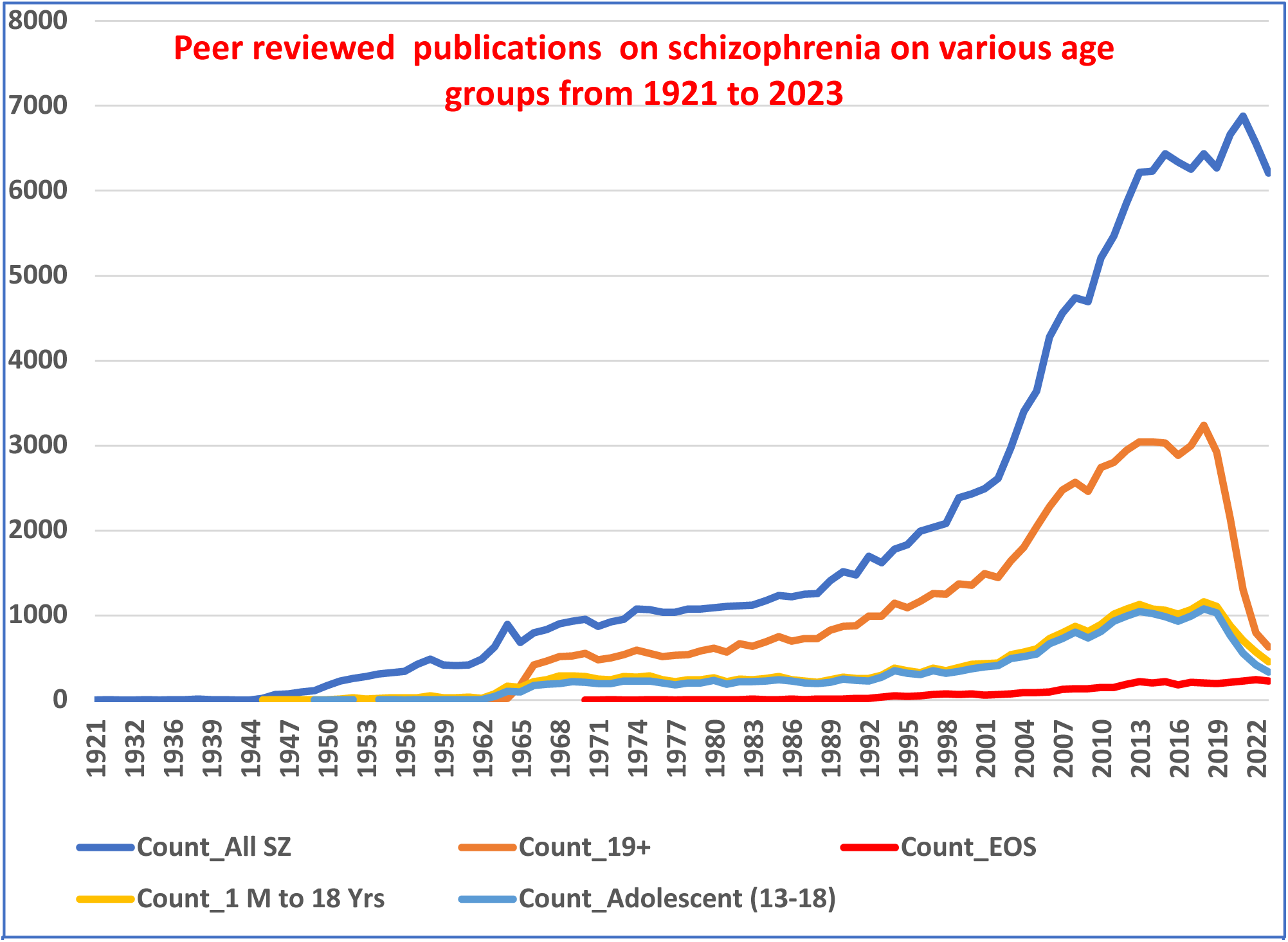
Peer reviewed publications from the PubMed on all schizophrenia and early onset schizophrenia. Dar blue line 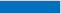 :All schizophrenia; Orange line 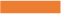: Schizophrenia in persons with age 19 and above; Light orange line 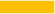: Publications on schizophrenia among persons 1 month old to 18 years; Light blue line 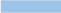: Publications on adolescents with schizophrenia between the ages of 13 to 18 years; Red line 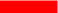: Publications with an explicit term “early-onset schizophrenia”

### Supplemental material 2

The implication of z-scoring is that roughly have the microstates are ‘on’ and half are ‘off’ over the course each fMRI acquisition, per node. Lowering this threshold is mathematically equivalent to using the fixation period mean activation as the threshold, for some unknown value less than zero, assuming the fixation period has lower mean activation than activation over the entire acquisition. In the opposite case, that the fixation period has higher mean activation, then some threshold value greater than zero will suffice. Regardless, the energy landscapes are more feature-rich at a threshold of zero. Additionally, the relationship between empirical probabilities of states and MEM-predicted probabilities is the smoothest at a threshold of zero. Thus, while other thresholds are acceptable, we examine only the threshold of mean activation for these reasons. As with the energy landscapes in the main manuscript, this analysis was performed only in the “example” model, not all permutations.

**Supplemental figure 2:**
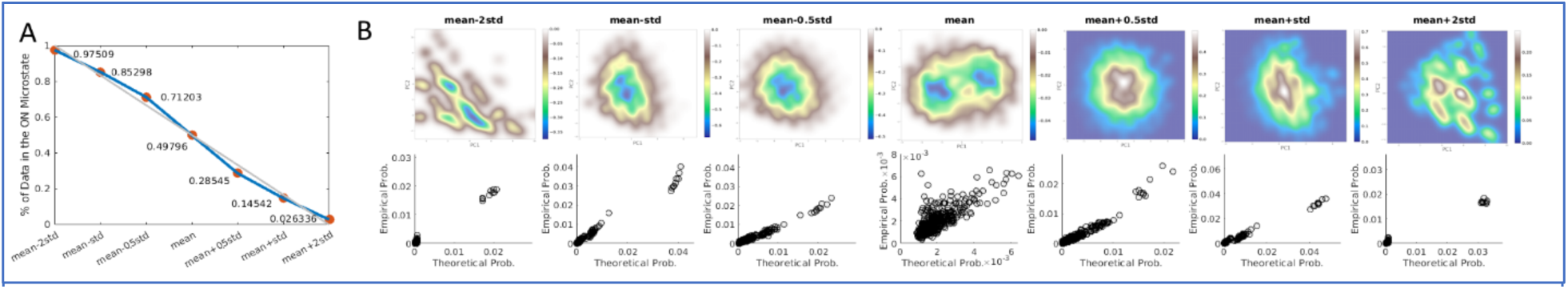
Threshold Effect on Energy. Left: The percent of data in the ‘on’ microstate across all subjects and timepoints after thresholding. Right: Seven thresholds were tested (x-axis), namely mean minus two standard deviations, mean minus one standard deviation, mean minus one-half standard deviation, mean, mean plus one-half standard deviation, mean plus one standard deviation, and mean plus two standard deviations. The middle threshold (mean) was chosen because in the threshold sweep results to the right, mean yielded the most complete “energy landscape” (top row, middle plot), and best fit of the empirical probabilities of state occurrence (bottom row, middle plot).

**<u>Supplemental material 3.</u> Maximum entropy model (MEM) fits.**

In our fitting algorithm, starting from some initial guesses, the values of *h_i_* and *J_ij_* in equation (1) were iteratively updated by comparing the empirically observed activation and co-activation frequencies (or equivalently mean activation level and covariance) of the imaged brain regions with those estimated from equation (2) and adjusting them with a learning rate, α = 0.2. Specifically, we calculated the estimated mean 〈σ_*j*_〉 and covariance 〈σ_*j*_σ_*l*_〉_*m*_ and performed the iterations of *h_i_* and *J_ij_* as follows:

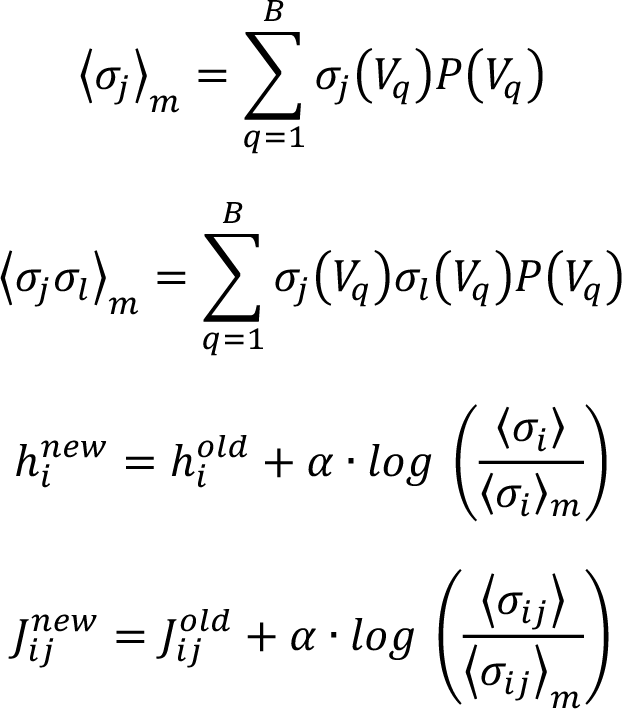

where ⟨σ_*i*_⟩ and 〈σ_*ij*_〉 are the empirically observed mean and covariance, respectively, for regions *i* and *j*. We ran 300 iterations of the gradient descent algorithm and that resulted in a good fit between the first-order (mean activity) and second-order (co-activation) statistics (see Section 2.6).

The MEM fitted to the combined data from control and AOS patients show a good fit to the first-order (mean activity) and second-order (covariance) statistics of the data.

**Supplemental Figure 3:**
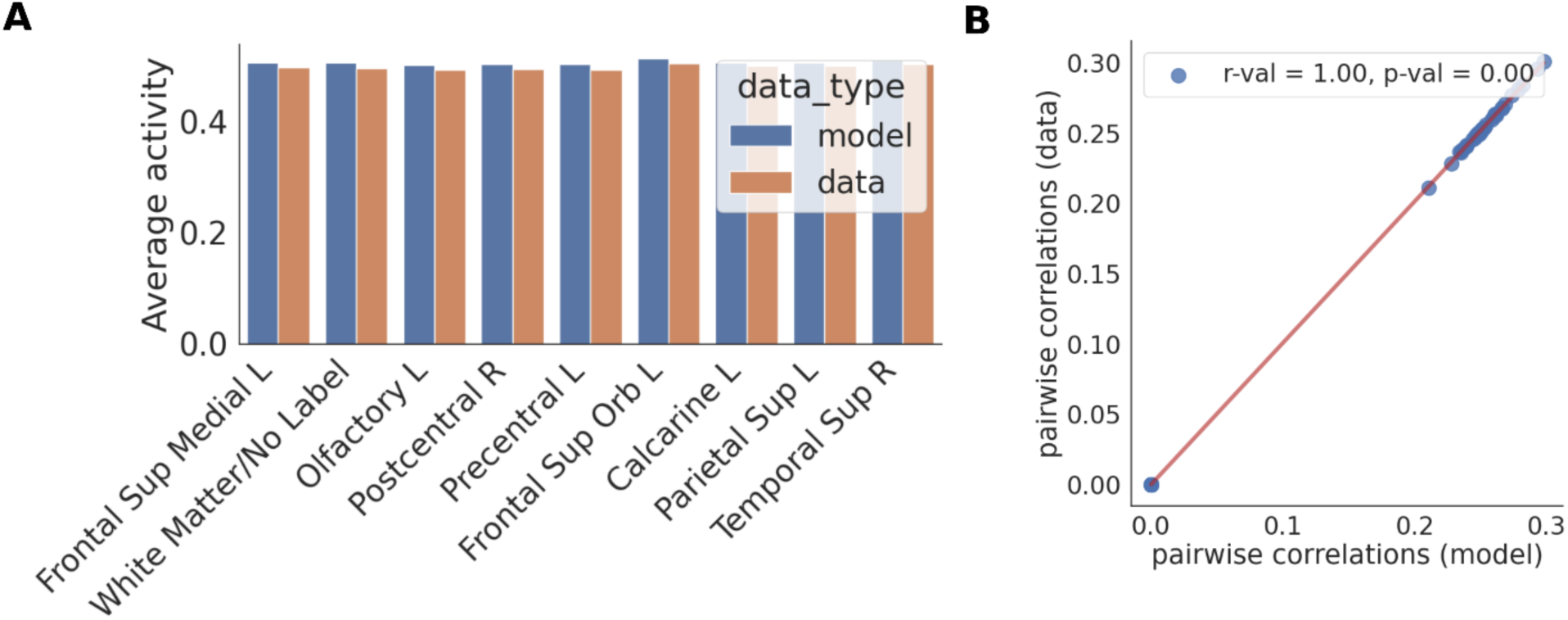
Goodness of fit of first-order and second-order statistics

**<u>Supplemental material 4.</u> Theoretical Versus Empirical Probability**

Comparing the predicted (theoretical) probability of state occurrence by the MEM to the energy of the same states shows a strict, monotonic relationship, as expected. While the MEM is internally tested for goodness of fit, this result confirms that the model is capturing differences in state occurrence frequency in terms of energy. It is a direct illustration of the relationship between energy and probability. Note that the empirical probabilities are what the model used to fit, so this result only indicates that the fitting succeeded.

**Supplementary Figure 4.**
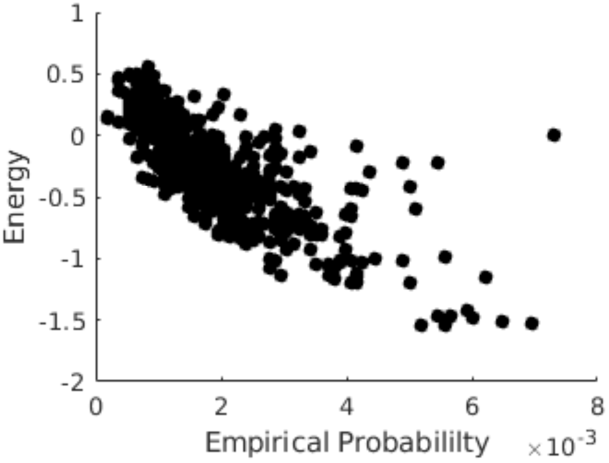
Top: Empirical probability of state occurrence is related to MEM-energy by state.

